# *Schistosoma mansoni* eggs modulate the timing of granuloma formation to promote transmission

**DOI:** 10.1101/2020.04.14.040626

**Authors:** Kevin K. Takaki, Gabriel Rinaldi, Matthew Berriman, Antonio J. Pagán, Lalita Ramakrishnan

## Abstract

Schistosome eggs provoke the formation of granulomas, organized immune aggregates, around them. For the host, the granulomatous response can be both protective and pathological. Granulomas are also postulated to facilitate egg extrusion through the gut lumen, a necessary step for parasite transmission. We used zebrafish larvae to visualize the granulomatous response to *Schistosoma mansoni* eggs and inert egg-sized beads. Mature eggs rapidly recruit macrophages, which form granulomas within days. Egg-sized inert beads also induce granulomas rapidly, through a foreign body response. Strikingly, immature eggs evade macrophage recruitment altogether, revealing that the eggshell is immunological inert. These findings suggest that the parasite modulates the timing of granuloma formation to its advantage, inhibiting foreign body granuloma formation until it reaches the optimal maturation and location for extrusion. At this point, the parasite secretes specific antigens through the eggshell to trigger granulomas that might facilitate egg extrusion.

## INTRODUCTION

Human schistosomiasis, caused by parasitic flatworms of the genus *Schistosoma*, affects more than 200 million people worldwide (WHO, 2019). Adult schistosomes live in the mesenteric venules of their definitive hosts, humans, where they produce eggs that are shed into the environment through feces or urine, depending on the species (Colley and Secor, 2014). Upon reaching fresh water, the eggs hatch, releasing the free swimming larvae (miracidia) that infect their intermediate snail hosts where they undergo asexual reproduction to release into the water the cercarial larvae that infect humans by penetrating the skin (Colley and Secor, 2014). In the case of *Schistosoma mansoni*, the most studied and geographically widespread species, the egg-laying adult pair resides in the mesenteric venous plexus. Upon maturation in the liver, the female and male adult worms pair up and migrate via the portal system to the mesenteric venules where they produce eggs (Nation et al., 2020). The eggs are shed by translocation through the venule and then the intestinal wall into the feces; however many become lodged in the intestinal wall or the liver (Hams et al., 2013; McManus et al., 2018; Nation et al., 2020; Schwartz and Fallon, 2018).

As the egg matures, it secretes antigens that provoke the formation of a granuloma - an organized aggregate of macrophages and other immune cells - around it (Ashton et al., 2001; Boros and Warren, 1970; Chiu and Chensue, 2002; Jurberg et al., 2009). For the host, the granuloma may play a dual function - both protective and pathogenic (Hams et al., 2013). On the one hand, it may protect the host by sequestering toxic egg antigens, and by preventing translocation of bacteria from the intestinal lumen into the tissues as the egg breaches the intestinal wall to exit the host (Costain et al., 2018; Hams et al., 2013; Pagan and Ramakrishnan, 2018; Schwartz and Fallon, 2018). On the other hand, the chronic granulomas around tissue-trapped eggs, particularly in the liver are the principal drivers of disease pathogenesis and morbidity (Hams et al., 2013; Pagan and Ramakrishnan, 2018). The chronic *Schistosoma* granuloma is complex in its cellularity with an abundance of myeloid cells, lymphocytes, eosinophils, and fibroblasts that act in concert to cause tissue pathology (Hams et al., 2013; Pagan and Ramakrishnan, 2018). The fibrogenic granulomatous response to the liver-trapped eggs causes damaging periportal fibrosis leading to portal hypertension and the development of esophogeal variaces which can rupture leading to internal bleeding and death (Colley and Secor, 2014; Pagan and Ramakrishnan, 2018).

While the granuloma’s role has mainly been studied from a host centric view, it has also been hypothesized that the early granuloma is critical for the parasite’s life cycle by facilitating the translocation of the eggs from the vasculature to the intestines and then into the feces for transmission to a new host (Dunne et al., 1983; Hams et al., 2013; Schwartz and Fallon, 2018). Because insights into the *Schistosoma* granuloma have been derived from single time point, histologic studies of human clinical samples and animal models-hamsters, mice and monkeys (Cheever et al., 2002; Hutchison, 1928), its role in translocation is understudied. The optical transparency of the zebrafish larva has enabled detailing of the early events of tuberculous granuloma formation in real-time using non-invasive, high resolution, serial intravital microscopy (Pagan and Ramakrishnan, 2018; Ramakrishnan, 2020; Takaki et al., 2013). Here, we have used the zebrafish larva to detail the events of early granuloma formation to *S. mansoni* eggs. We find that macrophage-dense epithelioid granulomas form rapidly around mature eggs. In striking contrast, we find that immature eggs are immunologically silent, failing to provoke even minimal macrophage recruitment. Given that inert beads induce epithelioid granulomas, this finding provides insight into how the egg might actively manipulate the timing of granuloma formation so as to prevent immune destruction or premature extrusion from the host. Our findings suggest a model where the eggshell inhibits the foreign body granuloma formation long enough for the miracidium to mature, at which point, antigens begin to be secreted through the eggshell to actively promote granuloma formation. This temporal control of granuloma formation may be critical in extruding the parasite exactly when it is ready to profit from the granuloma’s role in extruding the egg into the environment.

## RESULTS

### *S. mansoni* eggs induce epithelioid granuloma formation in the context of innate immunity

To study *Schistosoma* granulomas we used the zebrafish hindbrain ventricle (HBV), an epithelium-lined cavity to which phagocytes are recruited in response to chemokines and bacteria (Cambier et al., 2017; Cambier et al., 2014; Takaki et al., 2013; Yang et al., 2012)(Figure 1A). It has previously been shown that beads coated with *S. mansoni* soluble egg antigens (SEA) injected intravenously into mice get deposited in the lung where they induce macrophage recruitment and aggregation around them (Boros and Warren, 1971; Chiu et al., 2004). Using transgenic zebrafish with red fluorescent macrophages, we found that injection of SEA into the HBV induced macrophage recruitment within six hours (Figure 1B). Next, we implanted *S. mansoni* eggs into the HBV. Because the mature egg is relatively large (>50 μm diameter), we used a large bore borosilicate needle that allowed us to make an incision, grasp the egg and implant it into the HBV cavity in rapid succession (Figure S1, <u>Movie S1</u> and Methods, Figure 1C). Implantation of the eggs had no deleterious effect on larval survival; larvae implanted with either one or two eggs had a survival rate of 98%-100% at 5 days post-implantation (dpi), identical to the mock-implanted control group (n=50 per group). Implantation also did not change larval swimming behaviors or responses to tactile stimuli.

**Figure 1.**
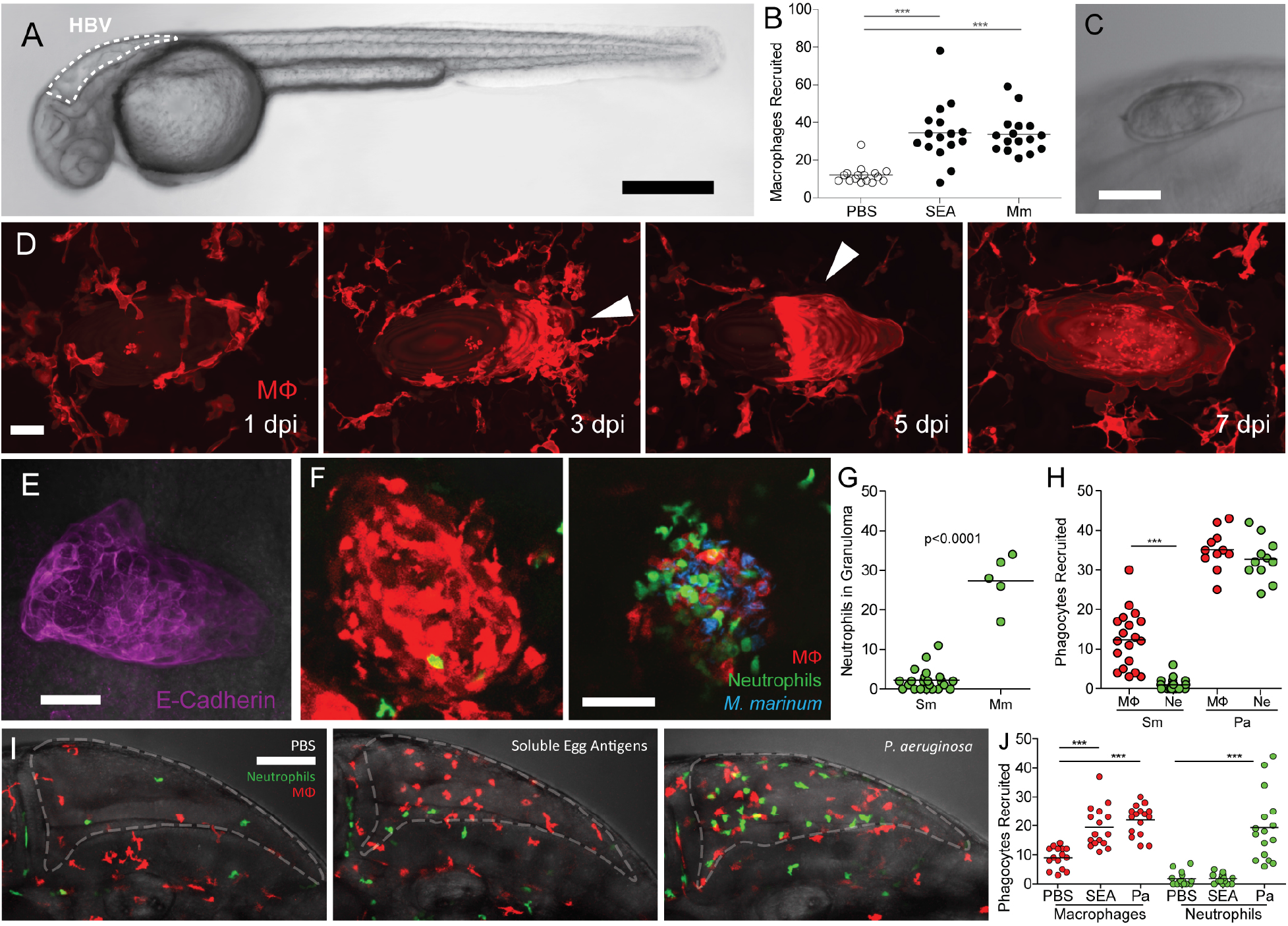
*S. mansoni* eggs induce epithelioid granuloma in larval zebrafish. (**A**) Zebrafish larvae at 30 hours post-fertilization with hindbrain ventricle (HBV) outlined. Scale bar, 300 μm. (**B**) Larvae were injected into the hindbrain ventricle with PBS, SEA or *Mycobacterium marinum* (Mm) and then assessed for macrophage recruitment at 3 hours post-injection. (**C**) Schistosome egg following implantation into the HBV. Scale bar, 75 μm. (**D**) Timelapse microscopy following the formation of the epithelioid schistosoma granuloma from 1-7 dpi. Arrowheads highlight the formation of the epithelioid granuloma with polar expansion. Scale bar, 25 μm. (**E**) Epithelioid granuloma immunostained using E-cadherin antibody. Scale bar, 50 μm. (**F** and **G**) Transgenic zebrafish larvae with red-fluorescent macrophages and green-fluorescent neutrophils were implanted with two *Schistosoma mansoni* (Sm) eggs each, or infected with 20 CFU *Mycobacterium marinum*, and then assessed at 5 dpi. (**F**) Confocal image of the schistosome egg granuloma (left) and *Mycobacterium marinum* granuloma (right), and (**G**) quantification of neutrophils. Scale bar, 50 μm. Data shown in (**H-J**) are from a single experiment each, (**F** and **G**), representative of two experiments, and (**B** and **E**) representative of four experiments. Horizontal lines in (B, G, H, and J) depict mean values. Statistics, (B, H, J) one-way ANOVA with (B) Dunnett’s or (H, J) Bonferroni’s post-test; G, two-tailed Student’s t test.

We assessed macrophage responses to the egg in transgenic zebrafish with red fluorescent macrophages. By one dpi, macrophages had arrived in response to the egg and were found flattened against it (Figure 1D; <u>Movie S2</u>). By 3 dpi, the macrophages had aggregated together on one part of the egg and this aggregate then progressively expanded to encapsulate the entire egg by 7 dpi (<u>Movie S2</u>). Even by 5 dpi, when the aggregate had not yet fully covered the egg, its macrophages appeared confluent with indistinct intercellular boundaries, suggesting they had undergone the characteristic epithelioid transformation associated with mature *Schistosoma* granulomas (Moore et al., 1977; Von Lichtenberg et al., 1973). Epithelioid transformation of the schistosome granuloma was confirmed by immunofluorescence staining for E-cadherin, the expression of which is its cardinal feature (Figure 1E; <u>Movie S2</u>)(Cronan et al., 2016; Pagan and Ramakrishnan, 2018).

It has been reported that *S. mansoni* eggs invoke macrophage-rich granulomas with very few neutrophils in contrast to *S. japonicum* eggs which recruit both macrophages and neutrophils (Chensue et al., 1995; Moore et al., 1977; Swartz et al., 2006; Von Lichtenberg et al., 1973). Likewise, we found that in the zebrafish, granulomas forming to *S. mansoni* eggs contained very few neutrophils (Figure 1F and 1G). In contrast, similarly-sized *Mycobacterium marinum* granulomas all contained neutrophils as expected (Figure 1F and 1G) (Yang et al., 2012). This pattern was established at the onset of egg implantation with macrophage but not neutrophil recruitment at 6 hours post-implantation (hpi), whereas the neutrophilic Gram-negative bacterium *Pseudomonas aeruginosa* recruited both types of cells, as expected (Figure 1H)(Yang et al., 2012). The lack of neutrophil recruitment has been attributed to the IL-8 neutralizing *S. mansoni* chemokine binding protein (smCKBP), more commonly known as alpha-1 (Smith et al., 2005), and accordingly the injection of SEA recruited only macrophages and not neutrophils, in contrast to *Pseudomonas aeruginosa* which recruited both (Figure 1I and 1J).

Next, we asked if the miracidium could survive within an epithelioid granuloma. We imaged individual eggs containing mature miracidia within organized granulomas at 5 dpi, and found they were still alive; the miracidium could be seen moving within the eggshell (Figure S2A; <u>Movie S3</u>). E-cadherin staining immediately after imaging confirmed that the granuloma macrophages had indeed undergone epithelioid transformation (Figure S2B). We also saw that in those cases where the eggshell had ruptured either during or after implantation, macrophages had entered into the eggshell and destroyed the miracidium, leaving the interior of the egg empty (Figure S2C; <u>Movie S3</u>). These findings were consistent with those in mammals showing that the intact eggshell protects the miracidium against destruction by host macrophages (Bunnag et al., 1986; Hutchison, 1928; Von Lichtenberg et al., 1973). Further confirming this, miracidia removed from the egg and implanted, rapidly recruited macrophages which destroyed them (Figure S2D).

In sum, we found that the key features of early mammalian responses to *S. mansoni* eggs are replicated in the zebrafish: selective macrophage recruitment to form bona fide epithelioid granulomas within days, which formed in the sole context of innate immunity. Our findings highlight that the miracidium tolerates granuloma formation as long as the eggshell is intact, a critical aspect of the *Schistosoma* life cycle that would depend on granulomas to enhance egg extrusion from the host. These granulomas most closely resemble intestinal granulomas in mice, which comprise mostly macrophages with fewer lymphocytes and eosinophils (Weinstock and Boros, 1983).

### Macrophages may be recruited to the egg by distinct consecutive signals

The serial observations in Figure 1D showing that the macrophages were rapidly enriched in one part of the egg, suggested that the first macrophages upon contact with the egg might be producing their own chemotactic signals that were stronger than those emanating from the egg. We pursued this idea by tracking macrophages by time-lapse microscopy in the first three hours after implantation. By 63 minutes, a small macrophage cluster consisting of 3-4 macrophages was already present on the egg (white arrowhead), and during the initial 15 minutes of the time-lapse, a new macrophage arrived and joined this macrophage cluster (Figure 2A). Meanwhile, three separate macrophages on a different part of the egg were found to come together into an aggregate (yellow arrowhead). By 180 minutes, additional macrophages had arrived at each of these clusters rather than to other parts of the egg. These results further corroborated that contact with the egg might render the first-arriving macrophages chemotactic, with the macrophages’ chemotactic signals being stronger than those of the egg.

**Figure 2.**
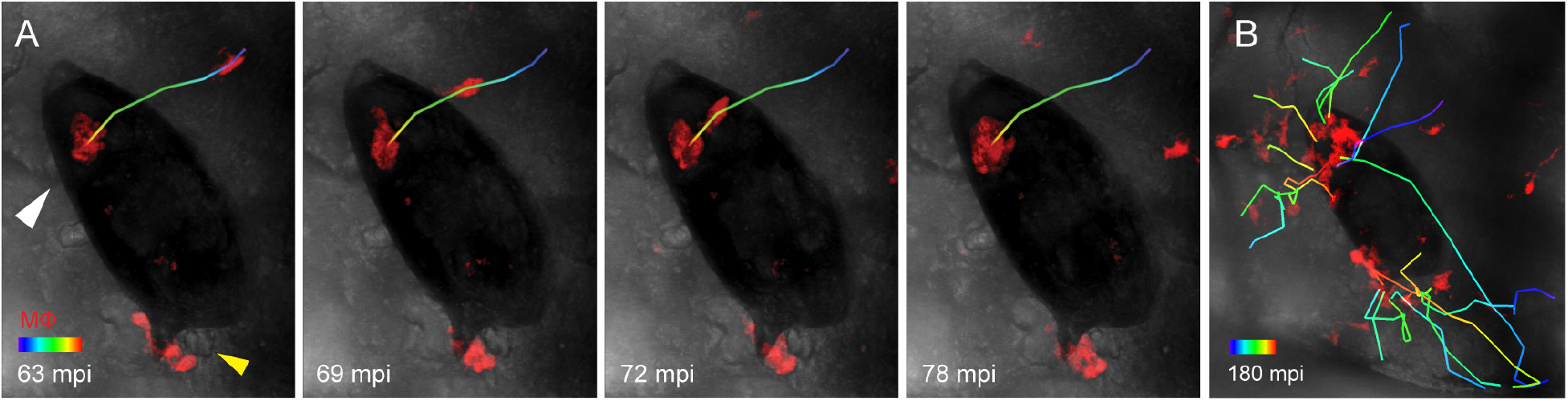
Dynamics of granuloma formation. (**A** and **B**) Timelapse microscopy tracking macrophage recruitment from 1-3 hpi. (**A**) Early aggregate of 3-4 flattened macrophages (white arrowhead) which recruits a newly arriving macrophage. Another aggregate (yellow arrowhead) condenses. (**B**) Macrophage tracks during the 2-hour timelapse from 1-3 hpi, showing the recruitment path to both macrophage aggregates. Scale bar colors represent time; later timepoints designated with longer wavelengths (red); scale bar length, 25 μm. Representative of two experiments.

### Immature *S. mansoni* eggs do not induce macrophage recruitment or granuloma formation

The egg matures six days after it is fertilized at which point it begins to secrete antigens (Ashton et al., 2001; Jurberg et al., 2009; Mann et al., 2011; Michaels and Prata, 1968). Accordingly, only viable, mature eggs are found to induce granulomas (Jurberg et al., 2009; Von Lichtenberg et al., 1973). We were able to sort immature and mature eggs based on their size and appearance (Figure S3A)(Jurberg et al., 2009). Granulomas formed around 25% of mature eggs (Figure 3A and 3B; Table S1). In contrast, no immature eggs invoked granulomas (Figure 3A and 3B). To corroborate this result, we used in vitro laid eggs at two days and six days post-fertilization (Figure S2A–S2C). Only the six-day eggs, and not the two-day eggs, induced granulomas (Figure 3C). These results were consistent with antigens secreted from the mature egg being the trigger for granuloma formation (Ashton et al., 2001; Boros and Warren, 1970; Chiu and Chensue, 2002). Heat-killed eggs produced fewer granulomas than live eggs, suggesting that a component of the granulomagenic egg antigens is heat-labile (Figure 3D)(Freedman and Ottesen, 1988; Klaver et al., 2015). Consistent with this, eggs that had been killed by storage at 4°C for 12 months so as to potentially inactivate all their antigens did not induce granulomas at all (Figure 3E).

**Figure 3.**
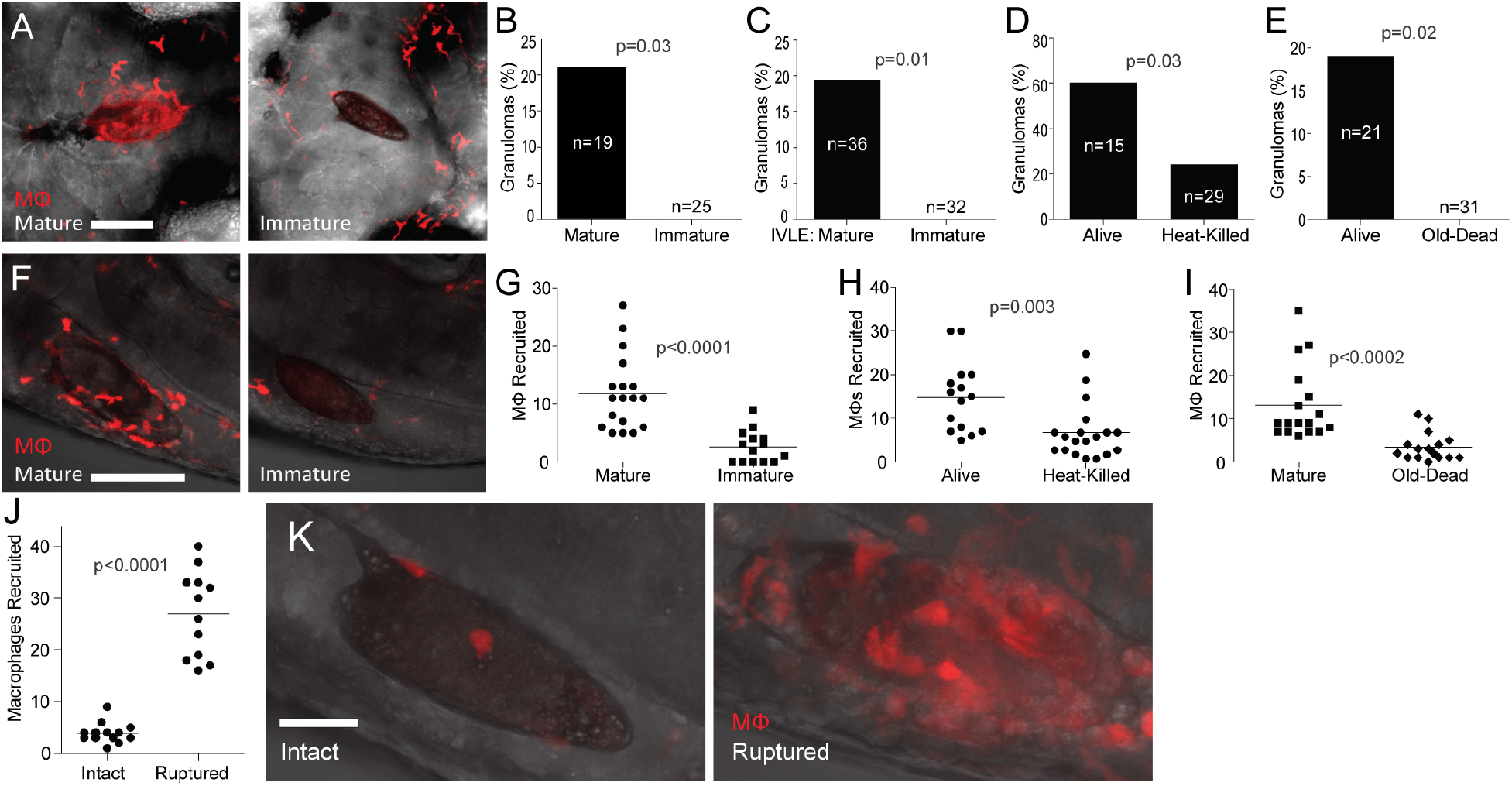
Immature eggs do not induce macrophage recruitment or granulomas. (**A**-**E**) Granuloma formation at 5 dpi comparing mature eggs with (**A** and **B**) immature eggs, (**C**) immature IVLE, (**D**) heat-killed eggs, or (**E**) old-dead eggs. Representative images in (**A**), scale bar 100 μm. Percent of animals with granulomas (**B**-**E**). (**F**-**I**) Macrophages recruited to, and in contact with mature eggs at 3 hpi in comparison to (**F** and **G**) immature eggs, (**H**) heat-killed eggs, and (**I**) old-dead eggs. Representative images in (**F**), scale bar, 100 μm. (**G**-**I**) Quantification of macrophages in contact with the implanted eggs. All experiments performed once, except for (**F**), which was a replicate of two experiments. (**J** and **K**) Immature eggs were implanted intact, or mechanically ruptured and then implanted. (**J**) Quantification and (**K**) representative images of macrophage recruitment at 6 hpi. Representative of two experiments. Scale bar, 25 μm. Statistics, Fisher’s exact test (**B**-**E**) and unpaired Student’s t-test (**G**-**J**).

We next asked if immature and dead eggs, while failing to form granulomas, could still induce an early transient macrophage recruitment. At six hours post implantation, immature or dead eggs had elicited Fewer macrophages than live mature eggs (Figure 3F–3I). Again, the immature and one year old eggs elicited fewer macrophages (23% and 28%, respectively, of macrophage recruitment to mature eggs) than the newly-killed eggs (40% of macrophage recruitment to mature eggs). These findings suggested that residual heat-stable antigens participate in the earliest stages of macrophage recruitment. Finally, we found that in contrast to intact immature eggs, ruptured immature eggs rapidly recruited macrophages (Figure 3J and 3K). Similar to the case with mature disrupted eggs, these macrophages entered the ruptured immature egg and killed the embryo (Figure 3K and data not shown). Together these results suggest that while the embryo and fully-mature miracidium have similar macrophage-recruiting properties, the intact egg at the two stages is fundamentally different in its ability to recruit macrophages. Our findings are consistent with the hypothesis that active antigen secretion from the mature egg is the stimulus required for the earliest macrophage recruitment step that is required to form granulomas.

### The immature *Schistosoma* egg evades foreign body granuloma formation

Our findings were consistent with macrophage recruitment occurring only in response to antigens secreted from the mature egg rather than to the eggshell itself. Granulomas form in response to inert foreign bodies (Pagan and Ramakrishnan, 2018), so why would the eggshell not induce a foreign body granuloma? We considered three possibilities. First, that it was too small to invoke a foreign body response; this seemed unlikely as very small inert particles, e.g. a tiny thorn, can provoke a robust foreign body response (Pagan and Ramakrishnan, 2018). Second, that the mechanisms to form foreign body granulomas were not yet operant in the developing zebrafish larvae; this too seemed unlikely given that the foreign body granuloma response is evolutionarily ancient, and epithelioid granulomas form in response to foreign bodies in invertebrates (Pagan and Ramakrishnan, 2018). Third, that the immature schistosome egg has specific mechanisms to evade foreign body granuloma formation. To distinguish among these possibilities, we implanted beads of three different chemically inert materials of the same size as the schistosome egg (Table S2). We chose Sepharose, which is hydrophilic, and polystyrene and polyethylene, which are hydrophobic. All induced macrophage recruitment within six hours (Figure 4A and 4B). By five days, epithelioid granulomas had surrounded the majority of the polystyrene and sepharose beads (Figure 4C–4E). The polyethylene beads were less granulomagenic with only 11% inducing bona fide granulomas (Figure 4C and 4D). However, even this weaker response was more robust than that of the immature eggs which did not even incite initial macrophage recruitment. We confirmed these findings with a head-on comparison of macrophage recruitment and granuloma formation in response to immature eggs or polystyrene beads in the same experiment (Table S2). Again, the polystyrene beads recruited macrophages by six hours and formed granulomas by five days, whereas the immature eggs did neither (Figure 4F and 4G). This result suggested that the immature egg specifically avoids being recognized as a foreign body. This could be because the immature egg secretes a specific product to inhibit macrophage recruitment, or that the eggshell is immunologically inert. To distinguish between these possibilities, we implanted an immature egg and a polystyrene bead adjacent to each other in the same animal. In every case, at six hours, macrophages were recruited only to the bead and not to the egg (Figure 4H and 4I). By 5 days post-implantation, granulomas had formed around the beads but none of the immature eggs (Figure 4J). These results supported the idea that the eggshell evolved to be immunologically inert so as to evade the ubiquitous foreign body macrophage recruitment and granuloma response.

**Figure 4.**
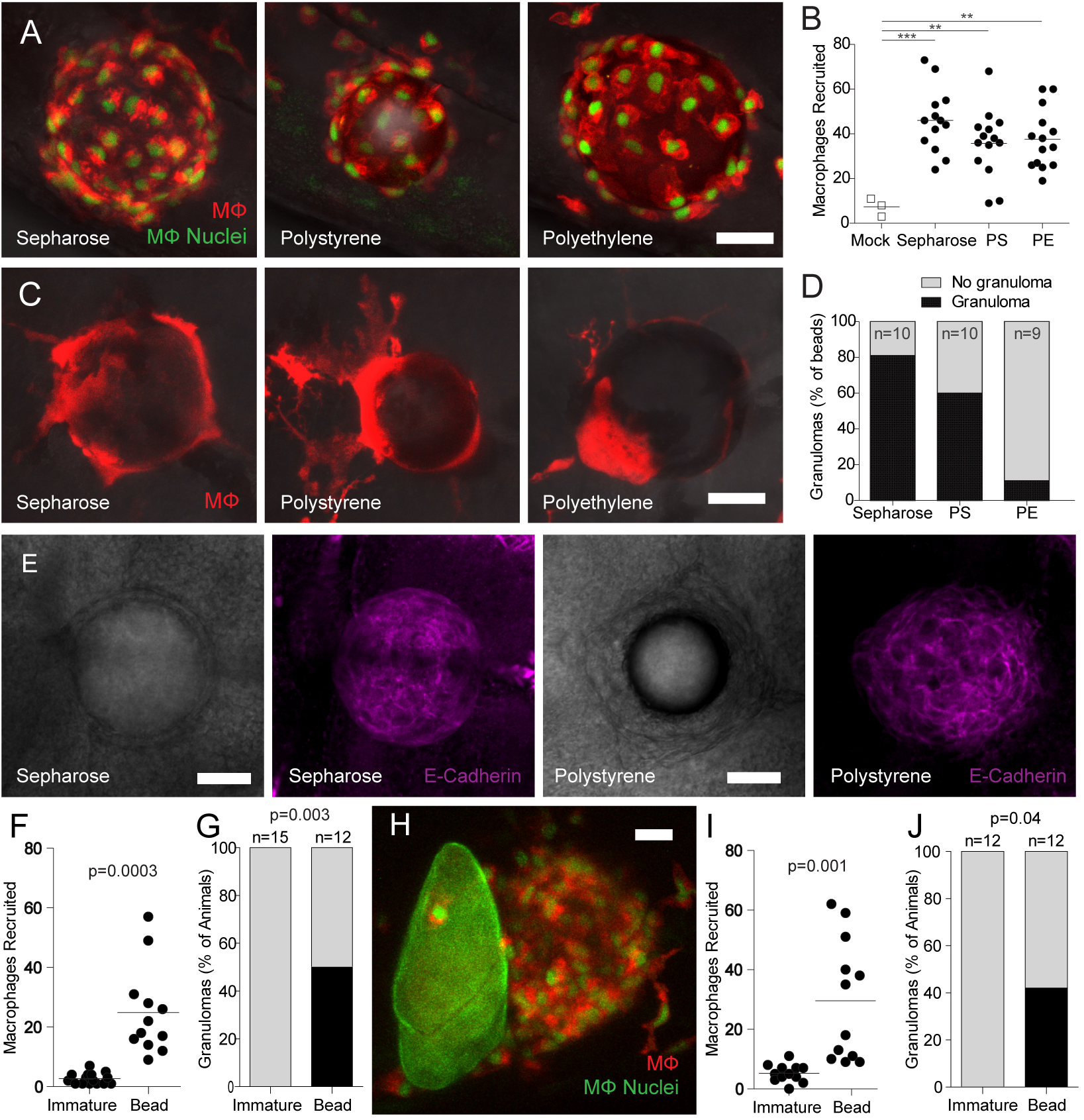
Chemically inert beads induce epithelioid granulomas. (**A** and **B**) Sepharose, polystyrene or polyethylene microspheres were implanted into transgenic zebrafish larvae carrying red-fluorescent macrophages with green nuclei and then assayed for macrophage recruitment at 6 hours post-implantation. (**A**) Representative images and (**B**) number of macrophages recruited per bead. (**C** and **D**) Microspheres were implanted into mpeg1:BB zebrafish larvae carrying the transgene for red-fluorescent macrophages and then analyzed for granuloma formation at 5 dpi. (**C**) Representative images and (**D**) quantification of granuloma formation. (**E**) Brightfield and fluorescence confocal microscopy of sepharose and polystyrene bead granulomas following staining for immunofluorescence against E-cadherin. (**F** and **G**) Larvae received a single immature egg or microsphere and were analyzed for (**F**) macrophage recruitment at 6 hpi or (**G**) granuloma formation at 5 dpi. (**H-J**) A single immature egg and polystyrene microsphere were implanted into the same larvae for analysis of macrophage recruitment and granuloma formation. (**H**) Representative image of an immature egg next to a microsphere, (**I**) quantification of macrophage recruitment at 6 hpi, and (**J**) quantification of granuloma formation at 5 dpi. Scale bars, 25 μm. Statistics, one-way ANOVA (**B**), unpaired Student’s t-test (**F** and **I**) and Fisher’s exact test (**G**-**J**). Replicates of one (**E** and **F-J**) and three (**A**-**D**) experiments.

## DISCUSSION

Research on *S. mansoni* granulomas has focused mainly on the organ-damaging fibrosis that ensues from granulomas forming around tissue-lodged eggs (Colley and Secor, 2014). Yet most *S. mansoni*-infected individuals are either asymptomatic or only mildly symptomatic (Hams et al., 2013), possibly because their granuloma or more response is more tempered. These individuals shed parasite eggs, highlighting that disease per se does not benefit parasite’s evolutionary survival. Rather, as in the case with many infectious diseases, human disease represents collateral damage the host-pathogen interaction, harming the host with little benefit to the pathogen (Relman et al., 2020). On the other hand, early, nuanced granuloma formation is thought to benefit both host and parasite for the same reason, expelling the parasite from the host (Dunne et al., 1983; Hams et al., 2013). While this idea is appreciated, it has been difficult to study extensively because of experimental limitations. Early or asymptomatic human infection seldom presents itself for study, and existing animal models are best suited to the study of late granuloma-associated pathology.

This work explores the earliest steps of Schistosoma granuloma formation that have not been captured in existing animal models. We show that as is the case with mycobacterial granulomas, bona fide epithelioid granulomas form in response to the Schistosoma egg in the sole context of innate immunity (Cronan et al., 2016; Davis et al., 2002). This should not be surprising given that epithelioid granulomas form in multiple invertebrate species in response to retained foreign bodies or even their own dead eggs (Pagan and Ramakrishnan, 2018). Yet, there has been at best a limited appreciation that adaptive immunity is not required for the formation of such an organized structure in the context of infectious granulomas, and indeed, the emphasis of schistosomiasis research on the late-stage granuloma has caused the focus to be on how the granuloma is modulated by adaptive immunity to become pathogenic (Hams et al., 2013; Pagan and Ramakrishnan, 2018). Given that Schistosoma eggs begin to be shed into the feces within days of being laid (deWalick et al., 2012), the transmission that occurs early in infection, at least, is likely promoted by these innate epithelioid granulomas. Our finding of the rapid epithelioid transformation of the granuloma also has relevance for granuloma-induced transmission later in infection when adaptive immunity is operant. The granuloma will still initially comprise only of macrophages (Pagan and Ramakrishnan, 2018), and it is this macrophage-dense granuloma that would be extruding the egg. Remarkably, intestinal granulomas, the ones that would extrude the eggs, are smaller than those in the liver, with a paucity of the lymphocytes and eosinophils that characterize liver granulomas (Weinstock and Boros, 1983). The rapid epithelioid transformation of the Schistosoma granuloma may help it to more efficiently extrude the eggs and hence propagate the parasite.

We have also gained understanding of the mechanics of early granuloma formation. Broadly speaking, granuloma formation in response to the mature egg proceeds in two discrete steps. In the first step, macrophages are attracted to secreted parasite antigens, and upon contact with the egg, appear to gain a chemotactic activity that outstrips that of the egg. This results in the subsequent macrophages being recruited to the existing macrophages forming a tight, aggregate that then pulls itself together to encapsulate the egg. It is noteworthy that epithelioid transformation precedes the complete covering of the egg, highlighting that this specialized macrophage transformation (Pagan and Ramakrishnan, 2018) constitutes an early response.

While these new details on how granulomas form around mature eggs are thought-provoking, more striking is the lack of even minimal macrophage recruitment by the immature egg. Given that like-sized beads recruit macrophages robustly and induce epithelioid granulomas, this finding provides new insight into Schistosoma biology. It reveals further nuance to the exploitation of the granuloma by the parasite. Not only must the parasite turn on granuloma formation through secretion of antigens, but it must also prevent the granuloma forming too soon. Premature granuloma formation may thwart granuloma for one of two reasons. The egg is laid into the blood stream, and needs to reach the wall of the blood vessel from which it extravasates and then penetrates the gut wall to be shed (deWalick et al., 2012). This process takes at least six days, perfectly synchronized with the time it takes for the miracidium to mature (Michaels and Prata, 1968; Pellegrino et al., 1962). Perhaps, premature granuloma formation might encumber its passage to the intestinal wall. Second, premature extrusion would remove the egg from the human tissue environment that is conducive to its maturation (Ashton et al., 2001).

Prior work has noted that the granuloma-inducing secreted Schistosoma antigens are secreted from the egg, rather than being incorporated into the eggshell, and that secretion occurs only after egg maturation (Ashton et al., 2001; Schwartz and Fallon, 2018). This work adds the key insight that the immunologically inert nature of the eggshell is a requisite counterpart of the Schistosoma transmission strategy. Our ability to directly compare granuloma formation around eggs and beads has been key to this insight. It will be interesting to determine how the eggshell remains immunologically inert in the context of granuloma formation, particularly so because eggshell proteins induce antibodies in humans (Dewalick et al., 2011; deWalick et al., 2012). Foreign body granuloma formation is a major complication of implanted devices (Pagan and Ramakrishnan, 2018). Identifying the chemical basis of the granuloma-silencing mechanism of the eggshell may have therapeutic implications to design inert materials for medical implants that alleviate this problem.

## AUTHOR CONTRIBUTIONS

A.J.P. and L.R. conceived the research project. K.K.T., A.J.P. and L.R. conceived and designed experiments and analyzed and interpreted data. K.K.T. performed the experiments. G.R. and M.B. provided knowledge, insights, experimental guidance and help with data analysis and interpretation. K.K.T and L.R. wrote the paper. K.K.T. made the figures. A.J.P., G.R. and M.B. edited the paper.

## ACKNOWLEDGMENTS

We thank S. Clare, C. Brandt, K. Harcourt, L. Seymour and C. McCarthy for assistance and technical support with animal infections and maintenance of the *S. mansoni* life cycle, P. Driguez and S. Buddenborg for support with *S. mansoni* egg preparation, G. Schramm for SEA preparations, and R. Keeble and N. Goodwin for zebrafish husbandry. This work was supported by Wellcome Trust core-funding support to the Wellcome Sanger Institute (award number 206194) (G.R., M.B.) and NIH MERIT award (R37 AI054503) and a Wellcome Trust Principal Research Fellowship (L.R.).

## SUPPLEMENTARY MATERIAL

### SUPPLEMENTARY FIGURES

**Figure S1.**
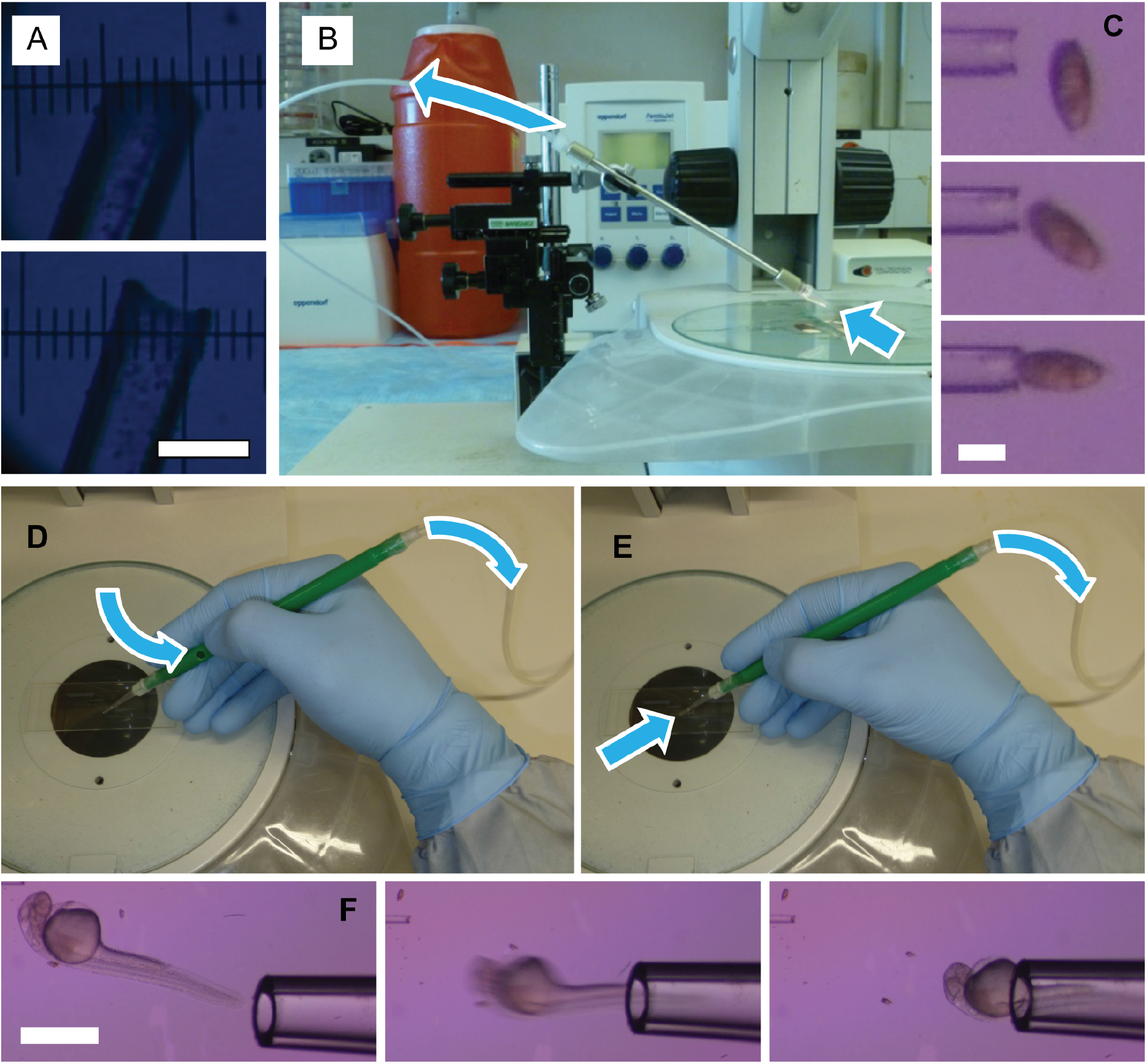
Implantation of schistosoma eggs into zebrafish larvae. (**A-C**) Capillary-Assisted Implantation Needle (CAIN). (**A**) Side and front profile of CAIN showing double-beveled point. Scale bar 50 μm. (**B**) CAIN attached to micromanipulator for X,Y, and Z control, as used by left hand of operator. Arrows indicated upward flow of fluid during grasping of egg. (**C**) Function of CAIN demonstrated by grasping *S. mansoni* egg. Scale bar, 50 μm. (**D-F**) Vacuum-Assisted MicroProbe (VAMP). (**D**) Occlusion of thumb hole re-routes aspiration pressure to tip (**E**), allowing for grasping of the larvae (**F**). Scale bar, 1000 μm. VAMP as described in Takaki, et al. 2013. Linked to Figure 1.

**Figure S2.**
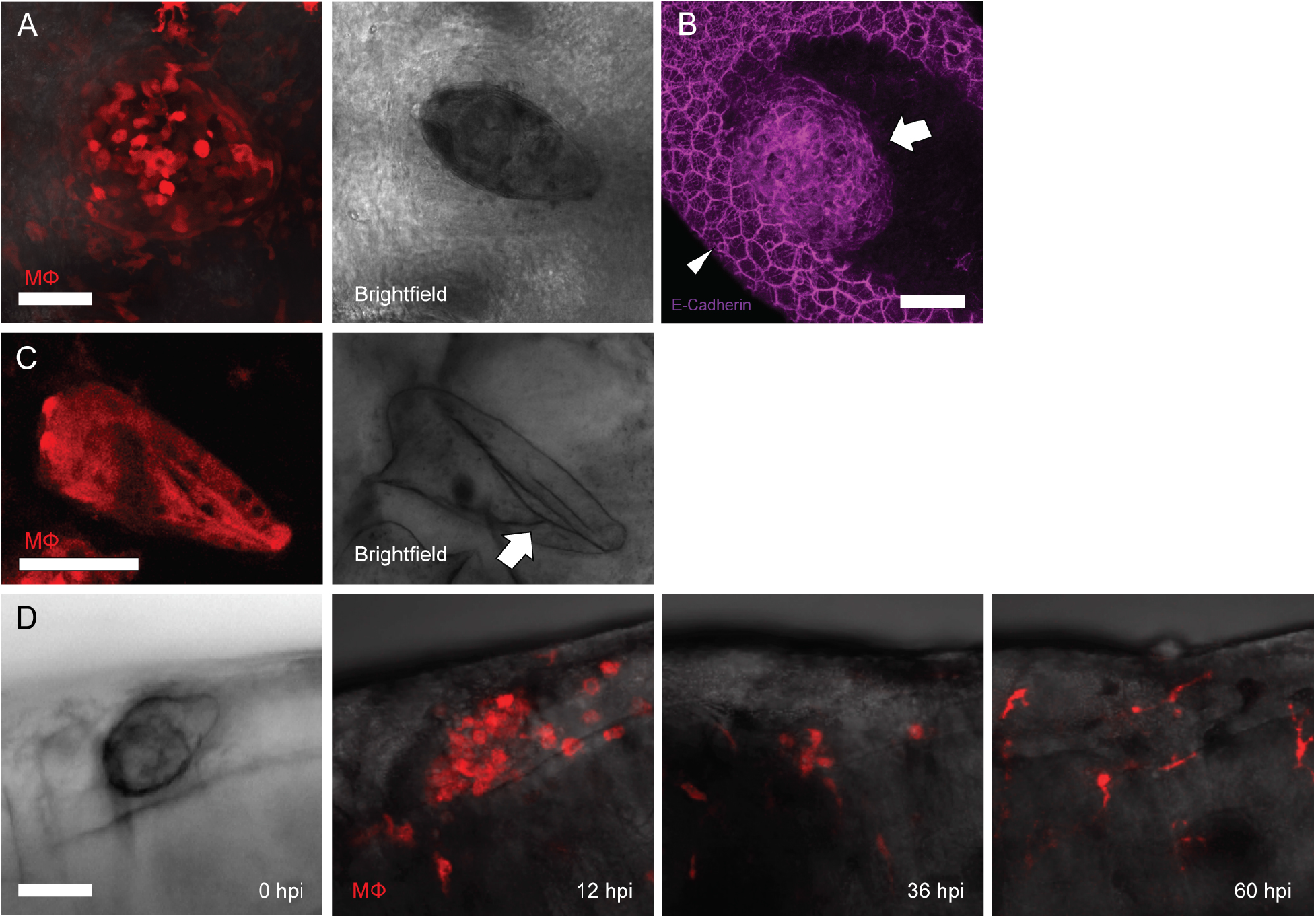
The eggshell protects the miracidium from being killed by host macrophages. (**A** and **B**) The parasite is alive within an epithelioid granuloma at 5 dpi. (**A**) Fluorescence and brightfield intravital microscopy. (**B**) Immunofluorescence staining of E-cadherin displaying maximum image projection (left), and optical sectioning (right). The outer-most stained structure is the epithelial lining of the hindbrain ventricle (arrowheads), and is not in contact with the epithelioid granuloma (arrow). (**C**) Fluorescence and brightfield microscopy of ruptured egg with macrophage infiltration resulting in degradation of parasite. Arrow, rupture of eggshell. (**D**) Brightfield and fluorescence timelapse microscopy of miracidia following implantation into the HBV. (**A**-**C**) observed routinely in experiments, (**D**) representative of two experiments. Scale bars, 50 μm. Linked to Figure 1.

**Figure S3.**
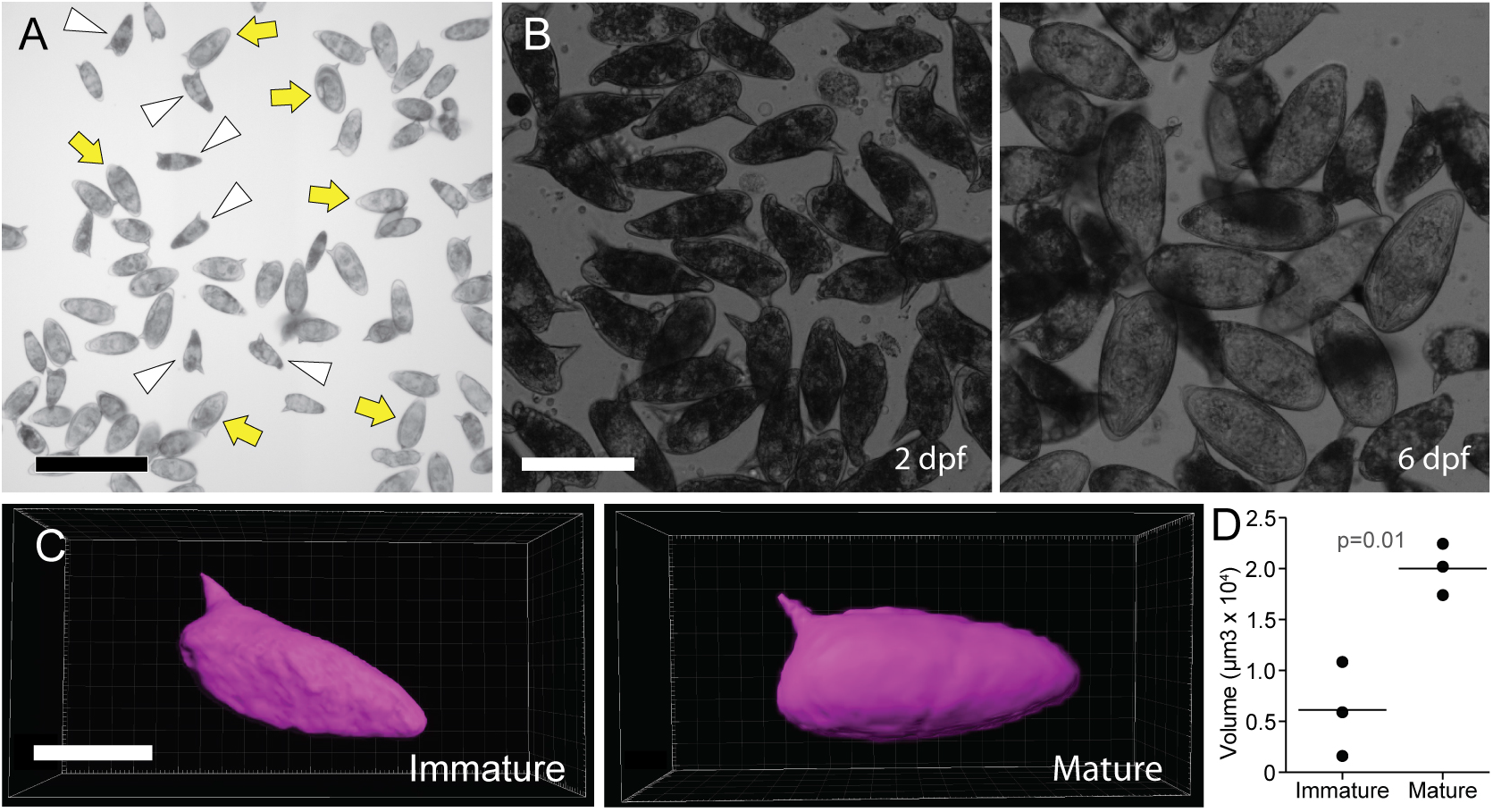
Sizes of mature and immature eggs. (**A**) *S. mansoni* eggs isolated from mouse livers. Immature and mature eggs, arrowheads and arrows, respectively. Scale bar, 300 μm. (**B**) Immature IVLE at 2 days post-fertilization (dpf), and mature IVLE at 6 days. Scale bar, 100 μm. (**C**) 3D rendering of Coomassie-stained eggs following confocal microscopy, and (**D**) volumetric analysis of three immature and mature eggs using 3D renderings shown in (**C**). Scale bar, 50 μm. Statistics, Student’s t test. Linked to Figure 3.

### SUPPLEMENTARY MOVIES

**<u>Movie S1</u>. Schistosome Egg Implantation**

Implantation of the schistosome egg using the CAIN and VAMP. Linked to Figure S1 and Methods.

**<u>Movie S2</u>. Formation of the Schistosome Egg Granuloma**

Timelapse 3D microscopy from 1-7 days post-implantation showing macrophage recruitment to the egg and granuloma formation around the egg. The last series of images shows E-cadherin staining around the same egg, done at the end of the time-lapse imaging. Linked to Figure 1.

**<u>Movie S3</u>. The parasite can withstand granuloma formation if the eggshell is intact**

The first series of images show a moving miracidium within an intact egg surrounded by an epithelioid granuloma five days post-implantation. The second series of images shows a ruptured egg which has been infiltrated by macrophages that are seen moving within the egg. Linked to Figure 1 and Figure S2.

**Table S1.** Granulomas form around some *S. mansoni* eggs

| Experiment | Animals implanted with egg | Eggs inducing granuloma | Eggs inducing granuloma (%) |
| --- | --- | --- | --- |
| 1 | 16 | 4 | 25 |
| 2 | 15 | 9 | 60 |
| 3 | 5 | 2 | 40 |
| 4 | 20 | 6 | 30 |
| 5 | 18 | 8 | 44 |
| 6 | 20 | 9 | 45 |
| 7 | 17 | 4 | 24 |
| 8 | 16 | 4 | 25 |
| 9 | 21 | 3 | 14 |
| 10 | 11 | 2 | 18 |
| 11 | 5 | 1 | 20 |
| 12 | 8 | 2 | 25 |
| 13 | 5 | 1 | 20 |
| 14 | 14 | 2 | 14 |
| 15 | 13 | 1 | 8 |
| 16 | 8 | 1 | 13 |
| 17 | 14 | 3 | 21 |
| 18 | 12 | 2 | 17 |
| 19 | 7 | 0 | 0 |
| 20 | 10 | 1 | 10 |
| **Mean** | **12.75** | **3.25** | **25%** |
| SEM | 1.2 | 0.6 | 3.2 |

**Table S2.** Size of implanted materials

| Implanted Material | Diameter (median, $\mu\text{m}$) | Volume (median, $\mu\text{m}^3$) |
| --- | --- | --- |
| Mature Schistosome egg | --- | 200,000 |
| Immature Schistosome egg | --- | 60,000 |
| Sepharose Agarose beads | 65 | 146,346 |
| Polystyrene beads | 45 | 47,713 |
| Polyethylene beads (large) | 70 | 175,909 |

### STAR METHODS

#### CONTACT FOR REAGENT AND RESOURCE SHARING

Further information and requests for resources and reagents should be directed to and will be fulfilled by the Lead Contact, Lalita Ramakrishnan.

#### EXPERIMENTAL MODEL AND SUBJECT DETAILS

##### Ethics Statement

All animal experiments were conducted in compliance with guidelines from the UK Home Office and approved by the Wellcome Sanger Institute (WSI) Animal Welfare and Ethical Review Body (AWERB).

##### Zebrafish Husbandry

All zebrafish lines were maintained on a recirculating aquaculture system with a 14 hour light −10 hour dark cycle. Fish were fed dry food and brine shrimp twice a day. Zebrafish embryos were housed in fish water (reverse osmosis water containing 0.18 g/l Instant Ocean) at 28.5°C. Embryos were maintained in 0.25 μg/ml methylene blue from collection to 1 day post-fertilization (dpf). At 24 hours post-fertilization 0.003% PTU (1-phenyl-2-thiourea, Sigma) was added to prevent pigmentation.

##### Fish Lines

Experiments requiring larvae with red-fluorescent macrophages were performed using Tg(mpeg1:Brainbow)^w201^ (Pagan et al., 2015). For experiments requiring analysis of neutrophils, Tg(lyz:EGFP)^nz117^ (Hall et al., 2007) were crossed with Tg(mpeg1:Brainbow)^w201^ (Pagan et al., 2015) to produce larvae with green neutrophils and red macrophages. Experiments assessing early macrophage recruitment in response to beads or ruptured immature eggs utilized Tg(mfap4:nlsVenus-2A-tdTomato-CAAX)(A. Pagán, unpublished). All zebrafish lines were produced in an AB background, with the exception of Tg(mfap4:nlsVenus-2A-tdTomato-CAAX) which utilized a mixed AB/TLF background.

#### METHOD DETAILS

##### Isolation and manipulation of schistosome eggs

The complete life cycle of *Schistosoma mansoni* NMRI (Puerto Rican) strain is maintained at the WSI by breeding and infecting susceptible *Biomphalaria glabrata* snails, and mice. Schistosome eggs were harvested as previously described (Mann et al., 2010). Briefly, anesthetized Balb/c female mice were infected by tail submersion in water containing *S. mansoni* cercariae collected from experimentally-infected snails, and 6 weeks later euthanized by an overdose of Euthasol (sodium pentobarbital and sodium phenytoin, 40 mg per mouse) delivered by intraperitoneal injection. Mixed-sex adult worms were collected by portal perfusion, washed and maintained in culture for *in vitro* laid eggs (IVLE) collection (below). The mouse livers were removed after the portal perfusion, minced with a sterile razor blade in 1X PBS containing 200 U/ml penicillin, 200 μg/ml streptomycin and 500 ng/ml amphotericin B (i.e. 2% antibiotic-antimycotic-ThermoFisher Scientific), and incubated with 5% clostridial collagenase (Sigma) in 1X PBS with 2% antibiotic-antimycotic at 37°C with shaking for 16 hours. The digested liver tissue mixed with the schistosoma eggs was washed three times with 1X PBS with 2% antibiotic-antimycotic by centrifugation at 400 g for 5 min at room temperature and serially filtered through a sterile 250 μm and 150 μm sieve. The eggs were then separated from the liver tissue by a sucrose-based Percoll gradient and washed three times as above. The eggs were kept at 37°C, 5% CO_2_ in DMEM supplemented with 10% FBS and 2% antibiotic-antimycotic. All the procedures were performed in sterile conditions inside a biological safety cabinet.

*S. mansoni* IVLE were harvested as previously described (Mann et al., 2010; Rinaldi et al., 2012). Briefly, schistosome mixed-sex worms collected by portal perfusion were washed with sterile 1X PBS and 2% antibiotic-antimycotic, placed in 6-well plates and cultured in modified Basch’s medium (Mann et al., 2010) at 37°C, 5% CO_2._ Two days later, the eggs laid *in vitro* by the cultured worm pairs were collected from the bottom of the well. For experiments using immature IVLE, eggs were implanted into zebrafish larvae soon after collection, and for experiments using mature IVLE, eggs were cultured in modified Basch’s medium at 37°C, 5% CO_2_ for 6 days before being implanted into zebrafish larvae. For experiments using heat-killed eggs, the eggs were killed at 90°C for 15 minutes and incubated in 1 mL of modified Basch’s medium for 3 days to wash away residual egg antigens. Old dead eggs were created by stored at 4°C for >12 months and were verified as unviable based on lack of miracidial movement and hatching. For experiments using ruptured immature eggs, the CAIN was used to apply downwards pressure in combination with a sideways motion over the glass slide.

##### Implantation of Schistosoma Eggs

Capillary-Assisted Implantation Needles (CAIN) were created by pulling borosilicate thin wall with filament capillaries (GC100TF-10, Harvard Instruments) using a micropipette puller (Sutter Instruments, P-2000) with the following settings: Heat = 350, FIL = 4, VEL = 50, DEL = 225, PUL = 150. The tips of pulled needles were opened with jeweler’s forceps and then double-beveled using a MicroForge-Grinding Center (MFG-5, Harvard Instruments). Micromanipulation was achieved using a 3-axis micromanipulator (Narishige, M-152) with pressure control using a FemtoJet Express microinjection unit (Eppendorf). The VAMP (Vacuum-Assisted MicroProbe) was previously described (Takaki et al., 2013).

Larval zebrafish were anesthetized and implanted at 30 hpf in 0.252 g/L tricaine (Sigma, A5040) in a modified Schistosomula Wash medium (500 ml DMEM, 5 ml 1M HEPES and 2% antibiotic-antimycotic) to prevent egg hatching during implantation. Anesthetized larvae were grasped using the VAMP and an incision was made in the forebrain region using the CAIN. After making an incision, a single schistosome egg was picked up using the capillary action of the CAIN, and passed through the incision and deposited into the hindbrain ventricle (<u>Movie S1</u>).

##### Implantation of Beads

Zebrafish larvae were implanted with Sepharose (Sigma, C9142), polyethylene (Cospheric, CPMS-0.96 63-75um and CPMS-0.96 27-32um), and polystyrene (Generon, 07314-5) microspheres in fish water containing 0.252 g/L tricaine (Sigma, A5040) using the same technique as with schistosome egg implantations. Bead volumes calculated using the median radius (1/2 diameter) and formula for the volume of a sphere (v=4/3πr^3^). Egg volumes determined by 3D confocal microscopy.

##### Hindbrain ventricle Microinjections

Hindbrain ventricle injection of bacteria and soluble reagents were performed under anesthesia with 0.252 g/L tricaine (Sigma, A5040) using a microinjection needle supplied to a FemtoJet Express microinjection unit (Eppendorf), with larval manipulation performed using the VAMP (Takaki et al., 2013) and (<u>Video online</u>).

##### Bacterial Infections

Zebrafish larvae were infected with 20 CFU of *Mycobacterium marinum* M strain (ATCC #BAA-535) constitutively expressing EBFP2 (strain KT30, (Takaki et al., 2013) or 200 CFU of *Pseudomonas aeruginosa* (strain MPAO1, courtesy of Professor Gordon Dougan). All bacterial procedures were performed using dedicated equipment separate from *Schistosoma* procedures, and disinfected with 70% EtOH after use.

##### Soluble Egg Antigens (SEA)

SEA was prepared by Dr Gabriele Schramm. Briefly, eggs were isolated from *S. mansoni*-infected hamsters as previously described (Schramm et al., 2018), and then homogenized in PBS, pH 7.5, using a sterile glass homogenizer. The homogenate was then centrifuged at 21 krcf for 20 minutes. Supernatants were pooled and then dialyzed overnight in PBS using a 3.5 kDa molecular weight cutoff dialyzer. Sample was then centrifuged at 21 krcf for 20 minutes, and supernatant (SEA) was aliquoted and stored at −80°C. SEA was quantified for protein concentration using the Micro-BCA assay (Pierce, 23225), and quality controlled by SDS-PAGE and western blotting against the *S. mansoni* antigens, Omega-1, Alpha-1, and Kappa-5. Quality control for low LPS content was performed using the Chromo-LAL assay (Associate of Cape Cod, Inc., C0031-5). SEA was injected at 2 ng per hindbrain ventricle (1.5 nL injection of 1.4 mg/mL SEA).

##### Immunofluorescence staining

Immunofluorescence was performed as previously described (Cronan et al., 2016). Briefly, zebrafish larvae were fixed in Dent’s fixative overnight at 4°C, rehydrated in PBS containing 0.5% tween 20, and then blocked for 1 hour in PBDTxGs (PBS containing 1% BSA, 1% DMSO, 0.1% Triton X-100, 2% goat serum). Mouse anti-E-cadherin antibody, clone 36 (BD, 610181) was added at a 1/500 dilution followed by incubation overnight at 4°C. Larvae were washed in PBDTxGs and then Alexa Fluor 647 Goat Anti-Mouse IgG (H+L) antibody (ThermoFisher, A-21236) added at a 1/500 dilution followed by incubation overnight. Larvae were washed 5 times in PBDTxGs before analysis.

##### Confocal Microscopy

Zebrafish were anesthetized in fish water containing tricaine and then and mounted onto optical bottom plates (MatTek Corporation, P06G-1.5-20-F) in 1% low melting point agarose (Invitrogen, 16520-100) as previously described (Takaki et al., 2013). Microscopy was performed using a Nikon A1 confocal laser scanning confocal microscopy with a 20x Plan Apo 0.75 NA objective and a Galvano scanner, acquiring 30-80 μm z-stacks with 2-3 μm z-step intervals. Timelapse microscopy was performed at physiological temperature using a heat chamber set to 28°C (Okolab) with an acquisition interval of 2.5-3 minutes. For multi-day timelapse imaging, zebrafish larvae were carefully removed using jeweler’s forceps and returned to their standard housing (see husbandry) for imaging at later timepoints.

#### QUANTIFICATION AND STATISTICAL ANALYSIS

##### Phagocyte recruitment

For quantification of phagocyte recruitment, fluorescence confocal microscopy was performed, capturing z-stack images at the designated timepoint following implantation of eggs or beads, or the injection of soluble antigens or bacteria. Experimental groups were then blinded, and 3D rendering of confocal images were used to count the number of phagocytes in contact with the schistosome egg or bead, or the number of phagocytes within the hindbrain ventricle following injection of soluble antigens or bacteria.

##### Determination of egg volume

Schistosome eggs were stained with Coomassie InstantBlue dye (Sigma, ISB1L) and imaged by confocal microscopy with the 641 nm laser and CY5 HYQ filter, 590-650 nm excitation and 663-738 nm emission. Using Imaris X64 (Bitplane) 3D surface rendering of the eggs were then generated and used to calculate the egg volumes.

##### Macrophage tracking

Time-lapse confocal images were used to generate 3D surface rendering of macrophages which were tracked over time using Imaris X64 (Bitplane).

##### Statistical analysis

Statistical analyses were performed using Prism 5.01 (GraphPad Software), with each statistical test used specified in the corresponding figure legend. Post-test p-values are as follows: ns, not significant; * p < 0.05; ** p < 0.01; *** p < 0.001; **** p < 0.0001. Where the n value is given and not represented graphically in the figure, n represents the number of zebrafish used for each experimental group.

### ADDITIONAL RESOURCES

Materials and data will be available upon request.

#### KEY RESOURCES TABLE

| REAGENT or RESOURCE | SOURCE | IDENTIFIER |
| --- | --- | --- |
| **Antibodies** |  |  |
| Rabbit polyclonal antibody against L-plastin | Abcam | ab210099 |
| Alexa Fluor 555 Goat anti-Rabbit IgG (H + L) antibody | ThermoFisher | A-21428 |
| Mouse anti-E-cadherin antibody, clone 36 | Becton Dickinson | 610181 |
| Alexa Fluor 647 Goat Anti-Mouse IgG (H + L) antibody | ThermoFisher | A-21236 |
| **Bacterial and Virus Strains** |  |  |
| *Mycobacterium marinum* M strain/pMSP12:EBFP2 | Takaki et al., 2013 | KT30 |
| *Pseudomonas aeruginosa* MPAO1 strain | Prof Gordon Dougan | MPAO1 |
| **Biological Samples** |  |  |
| *Schistosoma mansoni* eggs Puerto Rican strain | Dr Gabriel Rinaldi | N/A |
| Soluble Egg Antigens | Dr Gabriele Schramm | N/A |
| **Chemicals, Peptides, and Recombinant Proteins** |  |  |
| Sepharose agarose microspheres | Sigma | C9142 |
| Polyethylene microspheres (CPMS-0.96 63-75 µm) | Cospheric | CPMS-63-75um |
| Polyethylene microspheres (CPMS-0.96 27-32 µm) | Cospheric | CPMS-27-32um |
| Polystyrene microspheres (45 µm) | Generon | 07314-5 |
| Tricaine | Sigma | A5040 |
| Instant Ocean | Spectrum Brands | Instant Ocean |
| PTU (1-phenyl-2-thiourea) | Sigma-Aldrich | A5040 |
| Normal goat serum (10%) | ThermoFisher | 50197Z |
| Triton X-100 | Sigma-Aldrich | T8787 |
| Bovine Serum Albumin (BSA) | Sigma-Aldrich | A7906 |
| Fetal Bovine Serum (FBS) | ThermoFisher | 10082147 |
| Clostridial collagenase | Sigma-Aldrich | C5138 |
| HEPES (1M) | ThermoFisher | 15630080 |
| DMEM | ThermoFisher | 11965092 |
| Modified Basch's Medium | Mann et al., 2010 | N/A |
| Dimethyl sulfoxide (DMSO) | Fisher Scientific | BP231-100 |
| Low Melting Point (LMP) Agarose | Invitrogen | 16520-100 |
| Micro-BCA assay | Pierce Biotech Inc. | 23225 |
| Optical bottom plates | MatTek Corporation | P06G-1.5-20-F |
| Antibiotic-Antimycotic (100X) | ThermoFisher | 15240062 |
| Euthatal Solution for Injection (200 mg/ml) | Dopharma B.V. | N/A |
| **Experimental Models: Organisms/Strains** |  |  |
| *Schistosoma mansoni* NMRI Puerto Rican strain | Dr Gabriel Rinaldi | N/A |
| *Biomphalaria glabrata* | Wellcome Sanger Institute | N/A |
| Balb/c female mice | Wellcome Sanger Institute | N/A |
| Zebrafish (*Danio rerio*): wild type AB strain | Zebrafish International Resource Center | ZDB-GENO-960809-7 |
| Zebrafish: *Tg(mpeg1:Brainbow)*<sup>w201</sup> | Pagán et al., 2015 | ZDB-FISH-151204-7 |
| Zebrafish: *Tg(lyz:EGFP)*<sup>nz117</sup> | Hall et al., 2007 | ZDB-TGCONSTRCT-071109-2 |
| Zebrafish: *Tg(mfap4:nlsVenus-2A-tdTomato-CAAX)* | (A. Pagán, unpub.) | N/A |
| **Oligonucleotides** |  |  |
| Recombinant DNA |  |  |
| Software and Algorithms |  |  |
| NIS-Elements 4 | Nikon | N/A |
| Imaris X64 | Bitplane | N/A |
| Prism 5.01 | GraphPad | N/A |
| Illustrator CS5 | Adobe | N/A |
| Other: |  |  |

## Notes

### Competing Interest Statement

The authors have declared no competing interest.

